# Species level resolution of female bladder microbiota from 16S rRNA amplicon sequencing

**DOI:** 10.1101/2020.10.27.358408

**Authors:** Carter Hoffman, Nazema Y Siddiqui, Ian Fields, W. Thomas Gregory, Holly Simon, Michael A. Mooney, Alan J. Wolfe, Lisa Karstens

## Abstract

The human bladder contains bacteria in the absence of infection. Interest in studying these bacteria and their association with bladder conditions is increasing. However, the chosen experimental method can limit the resolution of the taxonomy that can be assigned to the bacteria found in the bladder. 16S rRNA amplicon sequencing is commonly used to identify bacteria in urinary specimens, but is typically restricted to genus-level identification. Our primary aim was to determine if accurate species-level identification of bladder bacteria is possible using 16S rRNA amplicon sequencing. We evaluated the ability of different classification schemes, each consisting of combinations of a reference database, a 16S rRNA gene variable region and a taxonomic classification algorithm to correctly classify bladder bacteria. We show that species-level identification is possible, and that the reference database chosen is the most important component, followed by the 16S variable region sequenced.

**Importance:** Species-level information may deepen our understanding of associations between bladder microbiota and bladder conditions, such as lower urinary tract symptoms and urinary tract infections. The capability to identify bacterial species depends on large databases of sequences, algorithms that leverage statistics and available computer hardware, and knowledge of bacterial genetics and classification. Taken together, this is a daunting body of knowledge to become familiar with before the simple question of bacterial identity can be answered. Our results show the choice of taxonomic database and variable region of the 16S rRNA gene sequence makes species level identification possible. We also show this improvement can be achieved through the more careful application of existing methods and use of existing resources.

## Introduction

The human body provides a wide range of habitats, supporting a variety of microorganisms that include bacteria, archaea, viruses and fungi, collectively known as the human microbiome(1). Recent evidence from sequence-based and enhanced culturing techniques have revealed a population of microbes that exist in the bladder, even in the absence of infection(2–7). The discovery of the bladder microbiota (also known as the bladder urobiome) has led researchers to question how these microbes influence the health of the host. Studies have shown that altered urobiome diversity is associated with urinary tract infections (8–10), neurogenic bladder(7, 11, 12), bladder cancer(13, 14), and urgency urinary incontinence (UUI)(4, 15). These studies collectively provide evidence that the bladder urobiome, while previously overlooked, is clinically relevant and warrants further investigation.

To study the relationships between the bacteria found in the human bladder and health of the host, it is necessary to accurately identify bacteria in a rapid and large-scale manner. Reliable methods of determining the bacterial identity of an unknown bacterium include Matrix Assisted Laser Desorption/Ionization-time of flight (MALDI-TOF) analysis or whole genome sequencing (WGS) of purified colonies; both techniques permit species-level identification of bacteria(16). However, culturing specific bacterial species is time consuming and laborious. This limitation has been circumvented by adopting culture-independent methods of sequencing DNA directly from an environmental sample, such as shotgun metagenomic sequencing and targeted amplicon sequencing, the latter most commonly involving the 16S rRNA gene(17, 18). These culture-independent sequencing methods are an attractive strategy because they can more accurately reveal microbiota diversity by identifying bacteria that are difficult to grow in culture(19).

Targeted amplicon sequencing of the 16S rRNA gene is a common method for identifying bladder bacteria in a large-scale manner(20, 21). This is particularly true in urine where the low microbial biomass(9, 22) often results in lower bacterial DNA quantities than what is required for shotgun metagenomic sequencing(23). When performing targeted amplicon sequencing, DNA is extracted from all cells in a sample. Next, the polymerase chain reaction (PCR) is used to amplify a small segment of the bacterial genome, typically spanning one or more variable regions of the 16S rRNA gene. This amplified DNA segment is subject to high-throughput sequencing, such as Illumina MiSeq or HiSeq short read technology. Bioinformatics are used to process the resulting sequences and identify the taxonomy of the bacteria(24).

When identifying bacteria using targeted amplicon sequencing there are three important components (**Figure 1**). These components are: 1) a <u>database</u> of DNA sequences annotated with taxonomic information; 2) the <u>identifier</u>, or DNA sequence of the unknown bacterium; and 3) a <u>classifier</u>, which is the algorithm that compares the unknown sequence to those in a database until the closest match is found. These components work together as a *classification scheme*. One common classification scheme that can be implemented through the use of the mothur(25) or QIIME2(26, 27) bioinformatic pipelines uses the Silva database(28), the V4 region from the 16S rRNA gene sequence as the identifier(29), and the Naïve Bayes algorithm(27) as the classifier. A limitation of this and many other commonly used classification schemes is that the taxonomic resolution is usually constrained to the genus level. Recently, several new approaches to sequence processing and taxonomy assignment have become available (e.g. amplicon sequence variant algorithms such as DADA2(30), and taxonomic classifiers such as Bayesian Lowest Common Ancestor (BLCA)(31)), which may improve resolution to the species level.

**Figure 1:**
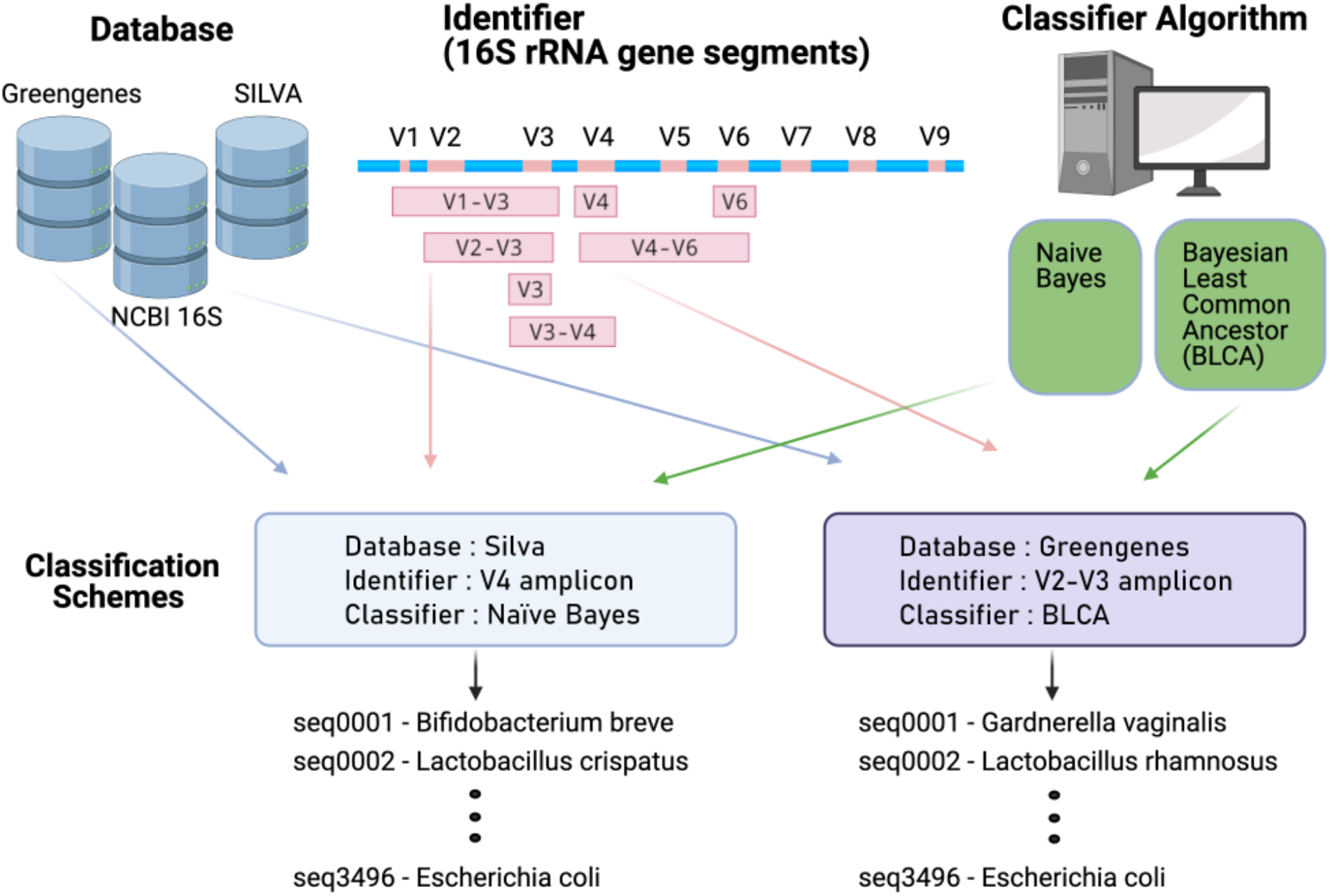
Model of the components that make up a classification scheme to assign taxonomy to unknown sequences. A classification scheme consists of a database, an identifier, and a classifier. The databases used in this study are the Greengenes, Silva, and NCBI 16S. The identifiers used in this study are subsequences of the 16S rRNA gene, computationally generated using published primers as coordinates on the gene sequence. These targeted amplicons are the V3, V4, and V6 variable regions of the gene, or span the V1-V3, V2-V3, V3-V4, and V4-V6 variable regions. The classifiers used in this study are the Naive Bayes and Bayesian Lowest Common Ancestor (BLCA) algorithms. One example of a classification scheme is the Silva database, V4 region identifier, and Naive Bayes classifier. Another example classification scheme is the Greengenes database, V1-V2 region identifier, and BLCA classifier. These two examples are distinct from each other and can have different outcomes when assigning taxonomy.

**Figure 1.**
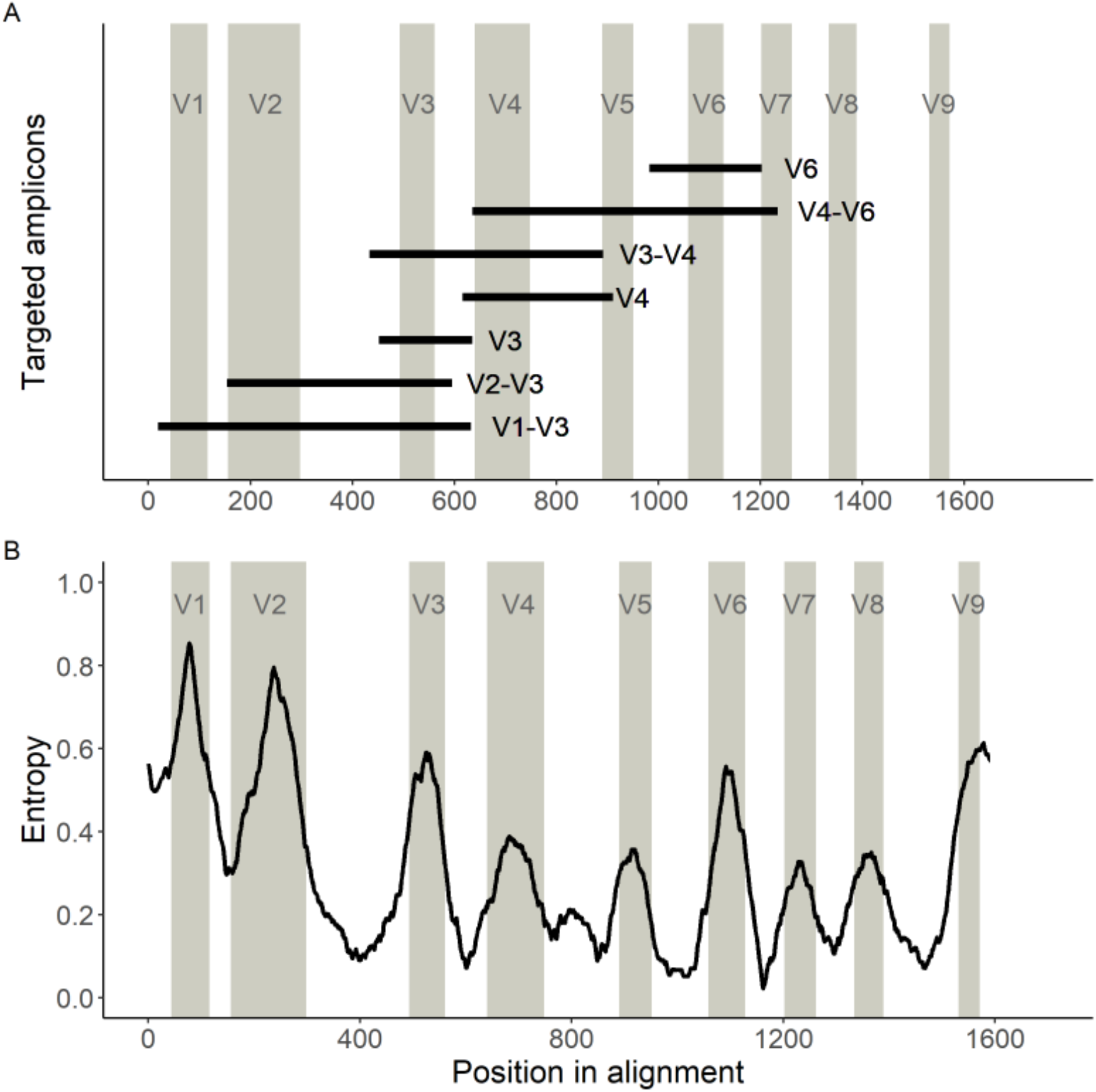
Variable regions of the 16S rRNA gene used in this study. A) Locations of the primers used in this study on the 16S rRNA gene. Locations of the predicted amplicons are shown as black bars in relation to the multi-sequence alignment (MSA) of the bacterial species described in Thomas-White et al. (2018). Gray columns are the locations of the known variable regions based on the sequence from *E. coli*. B) The information of variable regions, measured by entropy from a sliding window analysis of the MSA. Higher entropy indicates that the region has more variability across species, and therefore more information to identify a bacterial species. Lower entropy indicates that the region has little variability (i.e. is conserved) across species and therefore less information to identify a bacterial species.

Our primary aim was to determine if species-level identification of bladder bacteria is possible using 16S rRNA amplicon sequencing studies. To achieve this aim, we used known DNA sequences from a representative sample of bacteria found in the human female bladder, published by Thomas-White and colleagues(32). This dataset includes bacterial species that have been isolated from the bladder, cultured, and thoroughly characterized using WGS. We computationally tested multiple classification schemes to see which schemes identified these known DNA sequences at the species level. We evaluated several reference databases, variable regions (i.e. potential identifiers), and taxonomic classification algorithms for their ability to accurately identify bladder bacteria at the species level.

## Results

### Representation of bladder bacteria in 16S rRNA gene sequence databases

The Thomas-White genome sequencing dataset consists of 149 bladder bacterial isolates, representing 78 bacterial species from 36 genera(32). There are several databases available for bacterial identification using the 16S rRNA gene(33, 34). Of these, Greengenes (v.13_5)(35), Silva (v. 132)(36), and NCBI 16S Microbial (v. August 2019) were evaluated due to their widespread use in amplicon sequencing studies and availability of species-level annotation. At the genus level, all but one genus from the Thomas-White dataset (*Globicatella*) were present in the Greengenes database and all genera were present in the Silva and NCBI 16S Microbial databases. At the species level, all 78 bladder bacterial species were present in the Silva and NCBI 16S Microbial databases, whereas only 21 species were present in the Greengenes database.

### Information contained in variable regions differs for bladder bacterial species

Sequencing studies frequently focus on a small segment of the 16S rRNA gene that can be rapidly sequenced in a high throughput manner using short read sequencing technology, such as the Illumina HiSeq or MiSeq. To evaluate the performance of different variable regions as identifiers, amplicons were computationally generated from the Thomas-White genome sequencing dataset for the V1-V3, V2-V3, V3-V4, V4-V6, V3, V4, and V6 variable regions using published primers (see **Methods**). These computational amplicons were used to determine how well the currently available classification schemes can distinguish bladder bacterial species (**Figure 2A**). To assess different classification schemes, we tested multiple permutations of the variable regions listed above with different databases (i.e. Greengenes, Silva, or NCBI 16S Microbial) and different classifiers (i.e. Naive Bayes or BLCA, see **Figure 1**).

**Figure 2:**
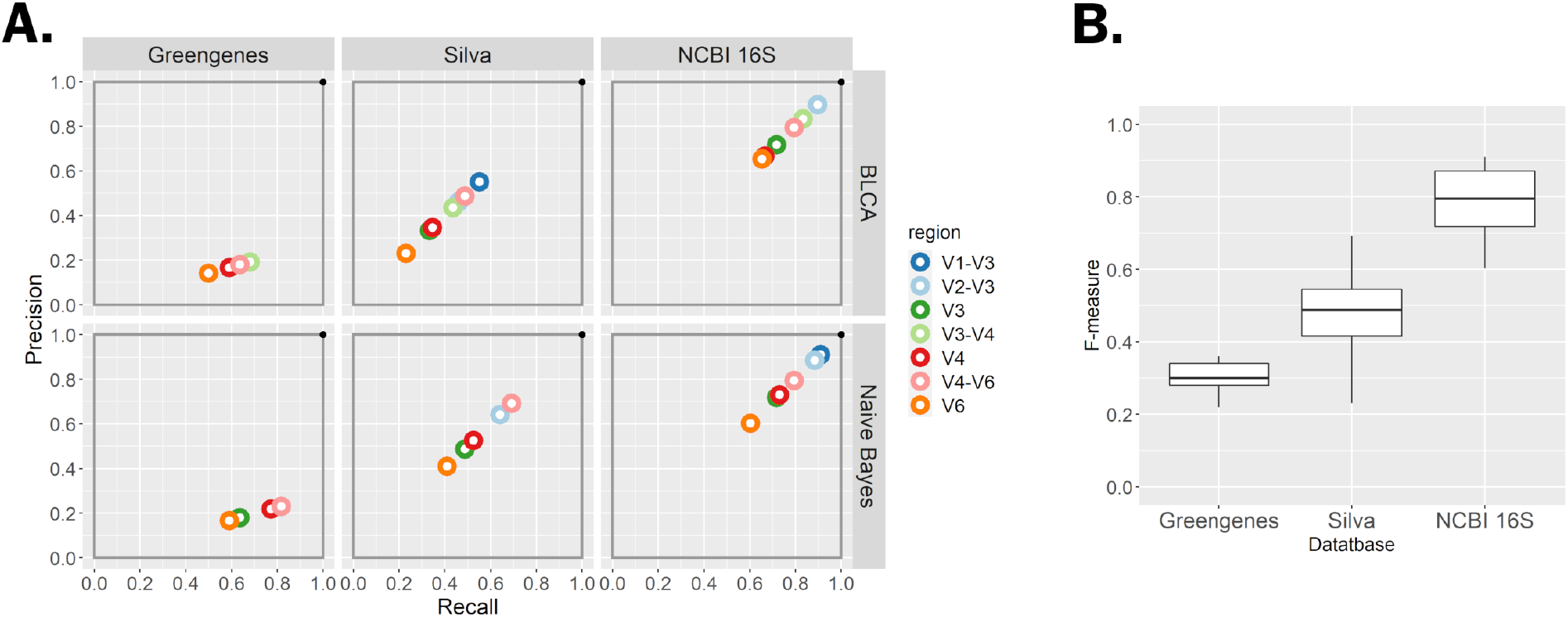
Overall classification scheme evaluation. **A.** The performance of each classification scheme is summarized by the precision (y axis) and recall (x axis) for each variable region (color). The best classification scheme would lie in the upper right-hand corner. **B.** Box plots depicting the median, quartiles, and range of the F-measure of all classification schemes by database. Overall, classification schemes using the NCBI 16S Microbial database had the highest F-measures and those using the Greengenes databases had the lowest.

To quantify the amount of information contained across variable regions of the 16S rRNA gene among commonly identified bladder bacteria, we performed a sliding window analysis on a multiple sequence alignment (MSA) of all 16S rRNA gene sequences from the Thomas-White dataset. We calculated entropy as a measure of information content along the MSA (**Figure 2B**). As expected, the defined variable regions contained regions of high entropy, suggesting variability across species, whereas variable regions were flanked by conserved regions with low entropy containing sequences that are similar among species. The V1 and V2 regions contained the highest entropy, while V7 and V8 contained the lowest.

### Evaluation of classification scheme performance

To evaluate the ability of currently available resources to identify bladder species, we calculated the recall, precision and F-measure for each classification scheme implemented (see **Methods**). For these calculations, we compared results from computational classification schemes to known bacterial species information from prior WGS. Each taxonomic classification was evaluated as a true match, true non-match, false match or false non-match based on whether the taxonomic classification was correctly assigned or not. Recall refers to the proportion of matches that the classification scheme correctly identified out of all possible matches. Precision refers to the proportion of matches that the classification scheme called correctly out of all classified matches. The F-measure is the equally weighted harmonic mean of recall and precision.

In general, classification schemes that use the NCBI 16S Microbial database perform the best as they have high recall and precision, thus the highest F-measures (median 0.79, range 0.60 – 0.91), regardless of which classifier is used (**Figure 3**). Classification schemes using the Silva database show reduced recall and precision, thus lower F-measures (median 0.48 range 0.23 – 0.69). Because the Greengenes database is missing many of the bacterial species found in the bladder, it is less precise. As such, classification schemes using the Greengenes database can have good recall but low precision resulting in a low F-measure (median 0.30, range 0.22 – 0.36).

**Figure 3:**
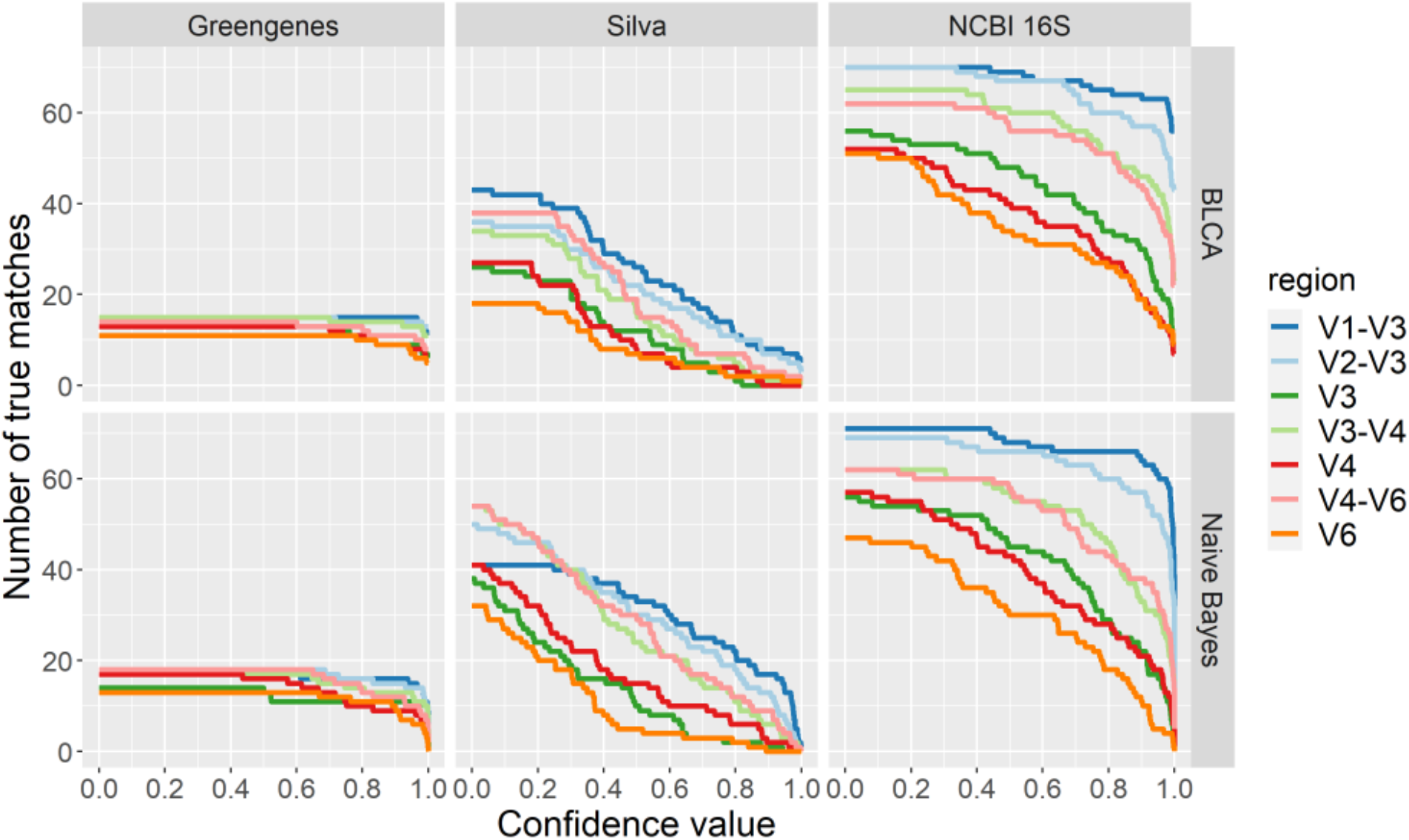
The number of true matches returned for each classification scheme across all confidence score values. As the confidence score value is increased where there is higher confidence of an accurate taxonomic assignment, the number of true matches dramatically decreases. For Greengenes and NCBI 16S databases, true matches especially decrease with confidence scores higher than 80%. With schemes using the Silva database, true matches steadily decrease with confidence scores as low as 30%.

While the choice of database appears to have the largest effect on the outcome of a classification scheme, the combination of identifiers and classifiers with specific databases show differences as well. For example, when using the Silva database, the classifier has an impact on the classification scheme outcome. With the Naive Bayes classifier, the identifiers yielding the highest F-measures are the V3-V4 (0.69) and V4-V6 (0.69) targeted amplicons. Using the BLCA classifier, the V1-V3 targeted amplicon has the highest F-measure (0.55).

### Confidence scores affect classification

The BLCA and Naïve Bayes classifiers used in this study will classify an unknown sequence even when the posterior probability for that taxon is very low. To account for this situation, a confidence score is calculated. This confidence score measures how much the classification changes through random permutation (bootstrapping) and the value reflects the “goodness of fit” of that classification. When lacking any knowledge of how to choose the best confidence score that minimizes the number of errors of a given classification scheme, using a predefined confidence score threshold is an option. We evaluated the performance of classification schemes against our test set when confidence score thresholds were used. For this analysis, matches that returned with confidence scores less than the designated confidence threshold were considered non-matches. **Figure 4** shows the effect of increasing the confidence score on the number of true matches returned by each classification scheme.

**Figure 4:**
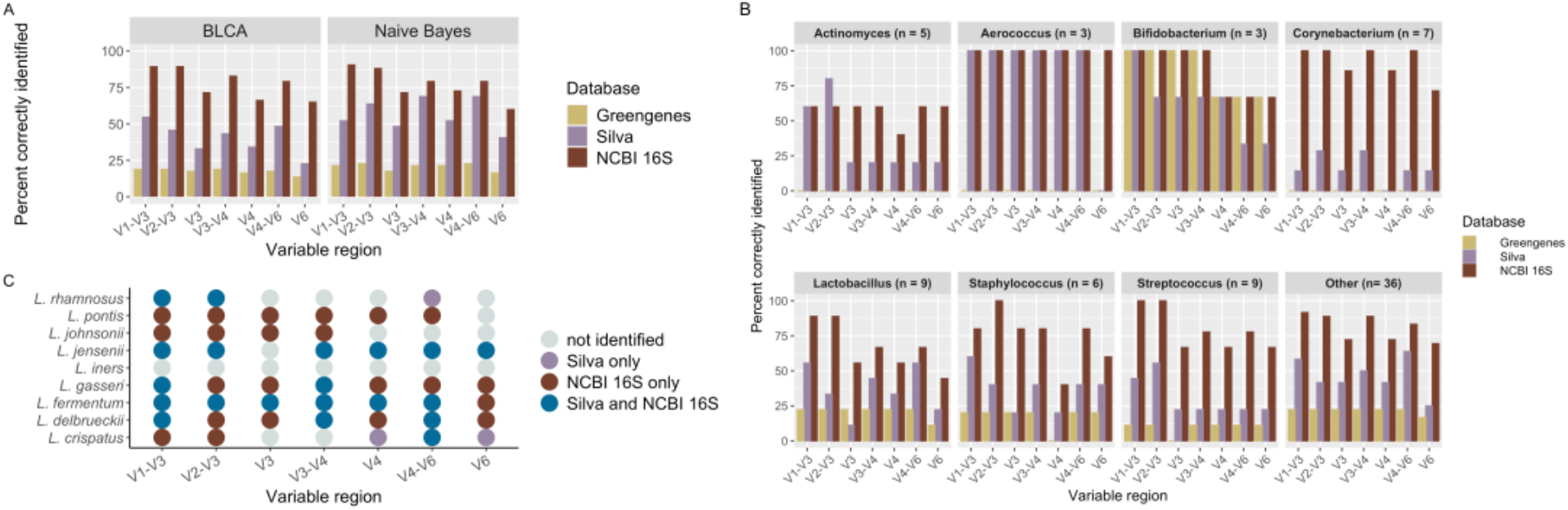
Percent of bladder bacteria correctly identified for each classification scheme. **A.** Overall results by classifier. With the commonly used V4 variable region and BLCA classifier, 17% of bladder bacteria are correctly identified using the Greengenes database, compared with 35% correctly identified using the Silva database and 67% using the NCBI 16S database. A similar trend is seen with the Naïve Bayes classifier. Using other variable regions can lead to improved species-level resolution to a maximum number of 91% correctly identified. **B.** The percent of bladder bacterial species identified with each database by variable region (BLCA). Genera with less than 3 species are grouped as Other. **C**. Lactobacillus species depends on the variable region and database used, with the NCBI 16S database having the most coverage, regardless of variable region chosen. The Greengenes database is not shown since it only classified two species (*L. pontis* and *L. delbrueckii*).

Almost all classification schemes had a decrease in recall when using a default confidence score of 80% (**Supplemental Figure 1**). This effect is especially marked for the classification schemes that use the Silva database, which shows an average 79.3% reduction in recall with higher confidence scores. Classification schemes that use the NCBI 16S database are unequally affected by increasing confidence scores; for example the V1-V3 identifier shows a slight reduction in recall (7.1% on average), while the V6 identifier shows the largest (43.3% on average). Classification schemes that use the Greengenes database are only slightly affected with different confidence scores resulting in smaller magnitude differences in recall.

Changes in the precision of classification schemes are mostly affected by the database used (**Supplemental Figure 1**), though confidence scores can further affect precision. For the classification schemes that use the NCBI 16S database, precision is generally improved with increasing confidence score, but at unequal amounts. For example, using the 80% threshold, the V1-V3 identifier shows a slight average increase of 3.5%, while the V6 identifier shows a large average increase of 39.7%. Classification schemes that use the Greengenes database show only minimal changes in precision. In contrast, classification schemes using the Silva database are differentially affected, with both reduction and gains in precision depending on the identifier and classifier combinations that are chosen. In general, for the Silva database, precision is reduced when using any 16S rRNA identifier with the BLCA classifier. However, a dramatic increase in precision is shown by the classification scheme composed of the Silva database, V4 identifier, and the Naive Bayes classifier. When ignoring a confidence score, this classification scheme has a precision of 52.6%, but increases to 85.7% when using a confidence score threshold of 80%.

The overall changes in how these classification schemes perform when using a 50% or 80% confidence score can be summarized by comparing the F-measure values shown in **Supplemental Figure 2**. In almost every classification scheme, the F-measure value decreases when a threshold is used. The classification schemes that use the Silva database clearly demonstrate this effect, which show a 66.9% reduction in F-measure values on average when higher confidence scores are used. The classification schemes that use the NCBI 16S database show slight decreases in the F-measure values, with the exceptions of those that use the V3, V4 and V6 regions as identifiers. Those classification schemes show a large 27.2% reduction in F-measure values on average. Finally, the classification schemes that use the Greengenes database have slight changes in their F-measure values, regardless of whether a threshold is used or not.

#### Amplicons spanning more than one variable region identify a higher number of bladder bacterial species

Amplicons spanning more than one variable region identified more unique bladder bacteria at the species level than amplicons spanning a single variable region. For example, with the commonly used V4 variable region and Naïve Bayes classifier, 21.8% of bladder bacteria are correctly identified with the Greengenes database, whereas 52.6% are identified with the Silva database and 73.1% with the NCBI 16S database (**Error! Reference source not found.**A). In contrast, using amplicons spanning more than one variable region such as the V1-V3 region, 91.0% of bacteria are correctly identified at the species level when using the NCBI 16S database.

#### Species identified depends on choice of database and variable region

While the results thus far have focused on summarizing overall performance of classification schemes for identifying bladder bacteria at the species level, we also sought to determine which classification schemes could be used to identify specific bacteria (**Figure 5B, Supplemental Figure 3**). Although the NCBI 16S database contains the largest representation of bladder species, some species were not identified with certain variable regions, if at all. For example, *Lactobacillus* species, which are commonly found in the female bladder and vagina (32, 37, 38), were overall best represented within the NCBI 16S database, with 8 out of 9 species being identified with the V1-V3 and V2-V3 variable regions (**Figure 5C**). However, the other variable regions only identified between 4 and 6 *Lactobacillus* species when using the NCBI 16S database. Interestingly, *L. crispatus* and *L. rhamnosus* were only detectable with specific combinations of variable regions and databases, and *L. iners* was not correctly identified from our dataset with any classification scheme.

Additionally, we found that there were important discrepancies for bacteria that are thought to play a role in bladder health and disease(39) (**Supplemental Figure 3**). Several bladder species, such as *Gardnerella vaginalis*, were only detected with the NCBI 16S and Silva databases. *Staphylococcus* species were poorly identified with the V4 region but were distinguishable with all other regions. *Streptococcus* and *Corynebacterium* species were best identified with NCBI 16S. *Escherichia coli*, a common uropathogen(40), is not well represented in any of the databases, and was only detected with the V4 region and the NCBI 16S database.

#### Validation of computational findings on V4 amplicon data

To evaluate the performance of our computational findings on actual data, we acquired targeted amplicon sequencing data from 24 urine samples. These urine samples were a subset of those that were used to derive cultures in the Thomas-White dataset and thus should contain the same bacteria. Sequencing data were generated as part of two other studies using Illumina sequencing of the V4 region of the 16S rRNA gene(4, 41). We reprocessed the raw sequencing data (see **Methods**) and performed taxonomic classification to assess the performance of our computational findings. Since 16S rRNA amplicon sequencing will detect many more bacteria than those identified even with enhanced culture, we restricted the evaluation to only the bacteria that grew in culture from a given sample (see **Supplemental Figure 4**). We used Cohen’s Kappa to assess the agreement between species assignment in the V4 computational dataset to the experimental V4 dataset (see **Methods**). All classification schemes had moderate agreement (**Table 1**).

**Table 1.**
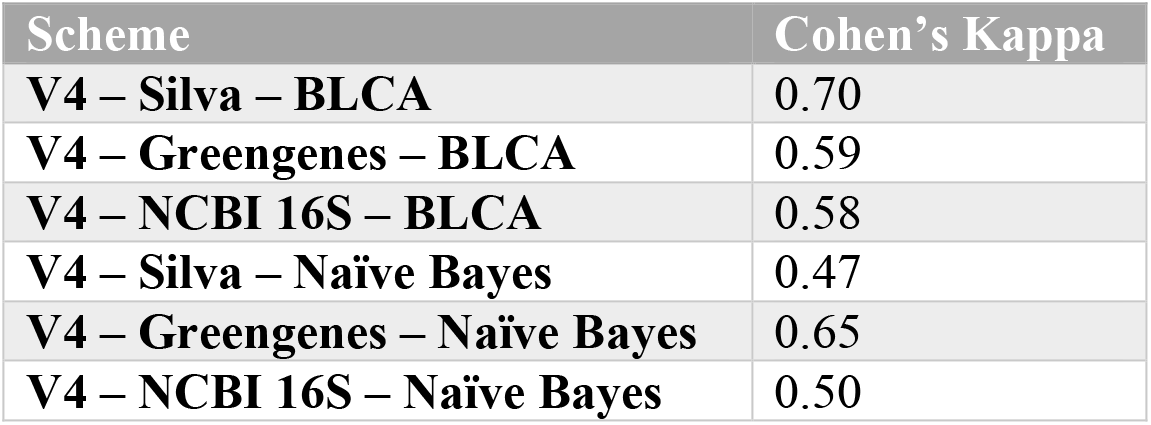
Cohen’s Kappa demonstrates moderate agreement of the computational and experimental amplicon datasets.

## Discussion

Our study demonstrates that it is possible to attain results at the species level with existing resources when performing targeted amplicon sequencing of urinary specimens. Though species-level taxonomic resolution is possible, it requires a carefully chosen classification scheme. Within the classification scheme, we found that the classifier chosen did not drastically affect the identification of species, but the database and identifier does. Importantly, we found that the reference database strongly influences the identification of bacteria at the species level. Overall, the NCBI 16S database performed the best, whereas the Greengenes database performed the worst using our approach.

While database selection is recognized as an important decision in taxonomy assignment for amplicon sequencing studies, few studies have compared the effect of different databases on taxonomy assignment. Furthermore, to our knowledge none have explored their use in identifying bacteria originating from the human bladder. Park and Won recently compared database performance on mock microbial communities and similarly found that the Silva database outperformed the Greengenes database(42). Several groups have also demonstrated that niche-specific databases perform the best for species level identification(43), an approach that has been taken in several microbiome research communities, such as the vaginal(44), oral(45) and gut(46).

For species level taxonomy assignments, the reference database must contain species-level information of the bacteria to be identified. We found that the Greengenes database does not currently contain many bacterial species that are found in the human bladder. In contrast, the NCBI 16S Microbial and Silva databases had representation of all species that were identified from prior studies of bladder bacteria. Thus, the latter two databases are better choices for evaluating bacterial species from the bladder. Even though both of these databases contain species of interest, the NCBI 16S database consistently outperformed the Silva database.

The classifiers used in this study are examples of two different strategies designed to overcome the common challenges of searching an extremely large dataset in order to find matching pairs of query sequences and reference records. While these two approaches are different in concept, we did not find significant differences in their performance for species-level classification of bladder bacteria. BLCA is an example of sequence comparison by pairwise alignment. The strength of this method is due to the fact that the similarities between two DNA samples are directly compared. This is a highly effective approach; however, until recent advances in computer technology, it remained impractical because of the computational burden. The Naive Bayes classifier is an example of a *k*-mer-based classification approach, and was designed to circumvent the computational challenges that are faced with use of a pairwise alignment classifier. However, there are limitations when using Naive Bayes for species-level identification. The first limitation arises from the database training process. If one taxon has more training examples than another, Naive Bayes generates unfavorable weights for the decision boundary(47). The second limitation is that all features (i.e. the *k*-mers generated from the DNA sequences) are assumed to be independent, and weights for taxa with strong dependencies among the associated *k*-mers are larger than those taxa with weakly dependent *k*-mers(47). A third limitation was that the available Naive Bayes Classifiers (such as those available through the RDP classifier(48) and QIIME2(27)) only had genus level information for the Silva and NCBI 16S databases. To use these for species-level assignment, it was necessary to retrain the classifier to use the specified databases at the species level. This training converts the DNA sequences to the calculated frequency that each *k*-mer occurs in a taxon.

Affordable sequencing of large-scale data is presently done with short read sequencing technology, such as Illumina MiSeq. This is currently limited to sequencing reads up to 300 nucleotides in length. Until full-length 16S rRNA gene sequencing can be achieved affordably on a large scale (e.g. technologies such as Oxford Nanopore and PacBio technologies), choosing the optimal region of the 16S rRNA gene for identification purposes remains a significant part of the experimental design. Thus, the variable regions that are used as identifiers require some consideration.

Our findings show that use of the V2-V3, and V1-V3 regions of the 16S rRNA gene allowed for the correct identification of the most bladder bacterial species when combined with the NCBI 16S database and either classifier. In general, amplicons that span more than one variable region perform better than those that contain single variable regions. This is likely due to the increased information available with longer reads. It is important to note that longer reads can have experiment limitations(49). While shorter variable regions, such as the V4 region, did not perform as well as longer amplicons, they were able to identify many bladder bacteria at the species level if the NCBI 16S database was used (52 out of 78). These shorter amplicons are widely used with Illumina sequencing and may be valid, depending on the study design and level of precision desired. However, other variable regions may be employed when more detailed information is desired.

This study is adds to a growing body of literature informing and improving experimental design for urobiome research. Several studies have now informed how the method of sample collection(2, 50), sample handling and storage(51), sample chemical composition(52), and DNA extraction protocol(52, 53) can influence downstream results and interpretations. By taking a computational approach to evaluating classification schemes, we were able to thoroughly assess the ability of various combinations of databases, identifiers, and classifiers to identify known bladder bacterial species. However, there are several practical limitations of amplicon sequencing that were not captured in this approach. We used published primers to create computational amplicons, but this may not reflect the efficiency of primers binding in actual experiments. Additionally, targeted amplicon sequencing generates a large number of overlapping reads and provides the data for methods to correct for errors introduced in the lab by the polymerase enzyme and through sequencing. Our method did not account for the experimental noise that is found in experimental data that may decrease the ability of the classification schemes to differentiate between bacterial species with similar sequences.

## Conclusion

Species level taxonomy assignment will greatly benefit studies focused on the urobiome and its relationship to bladder health and disease. Our results show that it is possible to reliably classify bladder bacterial species using targeted amplicon sequencing of the 16S rRNA gene variable regions with existing classification algorithms and databases. We determined that the most important component of the classification scheme is the database used, and that among those that were tested, the NCBI 16S database allows for the best identification of bladder species. Our validation with V4 amplicon data demonstrates that the predicted computational outcomes are a reasonable approximation for how a classification scheme will perform on real data. It can be expected that the alternate variable regions covered in this study, such as the V2-V3 region of the 16S rRNA gene, would have similar outcomes.

Importantly, we found that no single variable region gives 100% coverage of all bladder bacteria species. Thus, the choice of variable region may significantly affect the results of a given study. One approach to resolve this could be to use multiple amplicon sequencing or long read sequencing technology. These technologies are emerging and may prove to be beneficial for the urobiome community. Furthermore, no database has 100% coverage across a variable region. This could be resolved by using more than one database for classification, though this approach is complicated by differences in databases in terms of formatting, as well as conflicting classifications(54). Both of these components are important for planning experimental and computational aspects of urobiome studies, and should be considered when comparing results across studies.

## Material and Methods

### Code resources

All scripts that were written for this project can be found in the GitHub repository (https://github.com/KastensLab/BladderBacteriaSpecies). All scripts sourced from this repository are referred to as “custom.”

### The Thomas-White dataset

The 78 species of bladder bacteria used in this study were identified by culturing 149 bacterial isolates from 77 urine samples and performing whole-genome sequencing, as described in Thomas-White et al.(32). This set of identified species served as the basis for our computational analysis and is referred to as the Thomas-White dataset. For each species identified, the 16S rRNA gene sequence of the corresponding type strain was downloaded from the Silva v132 release (https://www.arb-silva.de/) on 4/27/2019. A *type strain* is the sequence of the cultured isolate that was subject to the metabolic, genotypic and phenotypic evaluations taken to define the bacterial species(55), and is the agreed bacterial organism to which the taxonomic name refers. Sequences were searched using the “[T]” filter setting, and sequences longer than 1450 nt with alignment and pintail quality scores greater than 95% were selected. For the species that had no hits, the taxonomic synonym (see below) was used as the search query, if available. One unidentified *Corynebacterium* species had no type strain available, and was excluded from the analysis.

### The V4 validation dataset

Targeted amplicon sequences from 24 urine samples, using the V4 region of the 16S rRNA gene sequence, is referred to as the V4 validation dataset. These 24 urine samples originated from a subsample of the women whose samples comprised the Thomas-White dataset. Sequencing data were generated as part of two other published studies using Illumina sequencing of the V4 region of the 16S rRNA gene(4, 41). The raw sequence reads were processed with DADA2 version 1.14.1(30) to generate amplicon sequence variants (ASVs). The ASVs were subjected to taxonomic classification with the BLCA algorithm.

### Synonyms of species

Species names have changed in response to advances in bacterial systematics. All currently known species synonyms were downloaded from the Prokaryotic Nomenclature Up-to-Date(56) (PNU) website on 1/5/2020. PNU includes information down to the strain level, but these entries were consolidated to the species level. For example, entries like *Enterobacter cloacae* and *Enterobacter cloacae dissolvens* are treated as synonyms of *Enterobacter cloacae*. Classification results were then checked for synonyms using the custom “validate_match_batch.py” script.

### Databases

The Greengenes database version 13_5 was downloaded on 9/23/19 from (http://greengenes.secondgenome.com/?prefix=downloads/greengenes_database/gg_13_5/). The Silva database version 132 was downloaded on 9/14/19 from (https://www.arb-silva.de/no_cache/download/archive/release_132/Exports/) as a FASTA formatted file. The NCBI 16SMicrobial database is bundled with the BLCA package, and is also available from (ftp://ftp.ncbi.nlm.nih.gov/blast/db/). For use with BLCA, each database was processed with provided scripts (`1.subset_db_gg.py` from BLCA for greengenes, <u>makeblastdb</u> utility from BLAST + suite combined with custom <u>write_taxonomy.py</u> for Silva, and `1.subset_db_acc.py` from BLCA for NCBI 16S Microbial). For use with Naïve Bayes, the FASTA file for each database was reformatted to work with QIIME2 using the custom “write_qiime_train_db.py” script. The Naive Bayes classifier was then trained with the “fit-classifier-naive-bayes” script QIIME2 script.

### Presence of Thomas-White species in databases

To verify that all species from the Thomas-White dataset were present in the databases used in this study, each database was first converted to a FASTA file (if needed) using the “blastdbcmd” utility included in the Blast+ suite. The FASTA file was then searched using regular expressions and the Linux command-line program *grep* for a match of each species in the dataset. The commands were implemented using the custom “species_in_db.bash” script. The presence or absence of each species was recorded.

### Multisequence alignment

The 16S gene sequences from the Thomas-White dataset were formed into a multi-sequence alignment using the T-coffee program(57). T-coffee version 12.00.7fb08c2 was downloaded from (http://tcoffee.org/Packages/Stable/Latest/) on 4/5/2019. Alignments were performed using the default settings.

### Sliding window analysis

Comparing the 16S rRNA gene sequences of the species in the Thomas-White dataset reveals regions of conserved sequence and regions of variability. The degree that variable regions of species differ from each other can aid the identification of each species; therefore, quantifying the amount of variability of a region across a set of species is important.

Sliding window analysis (SWA) is the method by which a list of subsequences are generated by taking successive groups of equal size, in the manner of a window of fixed length sliding across the full sequence. Quantifying the amount of variability along a MSA is achieved by combining SWA with calculating the Shannon Entropy contained in each column framed by the window.

The minimum Shannon entropy occurs when all nucleotides in a position (column) of the MSA are the same. The maximum occurs when all possible nucleotides in the MSA are present at that position. However, the Shannon Entropy treats gaps in a sequence as relevant, where in practice gaps reflect an absence of useful information. Multisequence alignments can generate many columns of gap characters due to insertions or deletions (indels) in the respective sequences that make up the MSA. A consequence of treating gaps as relevant is the Shannon Entropy will interpret these indel regions as conserved sequence. This limitation was overcome by weighting the entropy scores against gaps(58). The locations of known variable regions of the 16S gene sequence were validated, and the relative amount of variability was quantified, using the custom “weighted_ent.py” script.

### Primers

Amplicons were computationally generated from the Thomas-White genome sequencing dataset for the V1-V3, V2-V3, V3-V4, V4-V6, and V4 variable regions using published primers and the V3 and V6 regions using designed primers. The primer sequences used, listed in order of amplicon spanning variable region(s), forward primer name and sequence, reverse primer name and sequence are: **V1-V3**: A17F 5’-GTT TGA TCC TGG CTC AG-3’, 515R 5’-TTA CCG CGG CMG CSG GCA-3’(59, 60). **V2-V3**: 16S_BV2f 5’-AGT GGC GGA CGG GTG AGT AA-3’, HDA-2 5’-GTA TTA CCG CGG CTG CTG GCA C-3’(61, 62). **V3:** v3_579F 5’-THT TSS RCA ATG GRS GVA-3’, v3_779R 5’-GKN SCR AGC STT RHY CGG-3’. **V3-V4:** V3F 5’-CCT ACG GGA GGC AGC AG-3’, V4R 5’-GGA CTA CHV GGG TWT CTA AT-3’(63). **V4**: F515 5’-GTG CCA GCM GCC GCG GTA A-3’, R806 5’-CCT GAT GHV CCC AWA GAT TA-3’(18). **V4-V6:** 519F 5’-GTG CCA GCT GCC GCG GTA ATA-3’, 1114R 5’-GGG GTT GCG CTC GTT GC-3’(60). **V6:** v6_1183F 5’-CCG CCT GGG GAS TAC GVH-3’, v6_1410R 5’-AGT CCC RYA ACG AGC GCA-3’. Degenerate primer design was employed to generate primer sets for the V3 and V6 regions of the 16S rRNA gene that would anneal to as many species in the Thomas-White dataset as possible with DegePrime(64) (https://github.com/EnvGen/DEGEPRIME.git). DegePrime has the option to ignore columns of a MSA if the number of “-” characters exceed a user-defined threshold. The MSAs were preprocessed with this threshold set to .01. The main script of DegePrime was run using a degeneracy setting of 4096 and a window length of 18.

### Extracting computational amplicons

For each primer set, the DNA sequence bracketed by the forward and reverse primers was extracted from the multisequence alignment. Coordinates of the MSA were identified by searching the *E. coli* sequence (accession number EU014689.1.1541) included in the MSA for a match to the forward and reverse primer sequences, and then mapping those position to the MSA of the Thomas-White dataset. This procedure was done using the custom “extract_16s_vr.py” script and output as a multi-record FASTA formatted file.

### Taxonomic classifiers

Taxonomic classification was performed with Bayesian lowest common ancestor (BLCA) and Naïve Bayes classifiers. BLCA(31) was cloned from the GitHub repository https://github.com/qunfengdong/BLCA.git. For the 16S variable regions, the BLCA was run using default settings but pointing to the selected reference database, either Greengenes, Silva, or NCBI 16S. The Naïve Bayes classifier as implemented by QIIME2(27) was used with the Greengenes, Silva, and NCBI 16S databases and a confidence setting of 0, 50, and 80, but otherwise default settings.

### Evaluating computational results

To evaluate the taxonomic classification results of each classification scheme on the computational amplicon dataset, the custom “new_taxonomy_results_2020-3-14.Rmd” file was used. These scripts compare the results of each record pair (each comparison between the query sequence and sequence held in the reference database) from the classification scheme to the known identify of the query sequence from the Thomas-White dataset. All record pairs that were assigned a match by the classification schemes were evaluated according to the following definitions (**Figure 6**):

***True match*** - All record pairs assigned as a match that have identical genus and species labels.
***False match*** - All record pairs assigned as a match that did not have identical genus and species labels.
***False non-match*** - If a record representing a species in the Thomas-White dataset was present in the database, but was not assigned as a match, the record was evaluated as a false non-match.
***True non-match*** - All records in the reference database that were not in the Thomas-White dataset. While records assigned to this category were not used in evaluating the classification schemes in this manuscript, the definition is still included for completeness.

**Figure 6.**
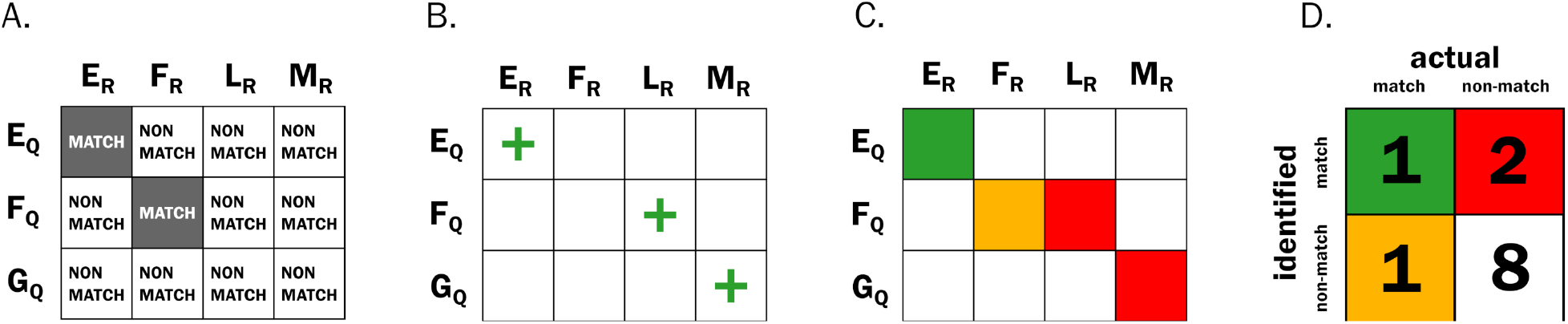
Example of classification evaluation used in this study. Suppose there is a classification scheme comprising a set of query sequences (the rows E,F,G) and the set of reference sequences (the columns E,F,L,M) held in a reference database. In this example, the number of reference records is greater than the query records, and the reference is missing a corresponding G record from the query set. **A**) If the query and reference record letters are the same, then they are designated as a **match**. If they are different they are designated as a **non-match**. **B**) Next, the classifier is allowed to assign record pairs as matches or non-matches for all query sequences, represented as green plus signs for matches and blank cells as non-matches. Some results are correct, and some are not. Note that despite the lack of a matching record in the reference database, the classifier still designated the (G:M) pair as a match. **C**) Using the definitions for assigning the classifications to the confusion matrix, there is one **true match** (green square), two **false matches** (red squares), one **false non-match** (yellow square), and 8 **true non-matches** (white squares). D) The cell values of the confusion matrix are then filled out, and performance measurements can be calculated. For this classification scheme, the precision is 1/(1+2)=.33, recall is 1/(1+1)=.5, and the F-measure is (2*.33*.5)/(.33+.5)=.40.

### Performance measures

Recall, precision and the F-measure were used to evaluate the performance of each classification scheme implemented. Recall refers to the proportion of matches that the classification scheme correctly identified (true matches) out of all possible matches (true matches plus false non-matches). Precision refers to the proportion of matches that the classification scheme called correctly (true matches) out of all classified matches (true matches plus false matches). The F-measure is the equally weighted harmonic mean of recall and precision. For this study, we chose to maximize recall over precision, because the number of true matches impacts the subsequent work on diversity measures, such as species richness and evenness(65).

### Evaluating V4 validation results

To determine the expected bacterial species in each sample, the results of the whole genome sequencing on the isolates cultured from the corresponding subject was used. The species of bacteria in the V4 sequencing data were identified using classification schemes composed of 1) the Greengenes, Silva, and NCBI 16S microbial databases 2) the V4 sequencing results as the identifier, and 3) the BLCA and Naive Bayes classifiers. For each classification scheme, the species identification from the computational and validation datasets was scored as a ‘1’ if correct, and ‘0’ otherwise. This resulted in a table for each classification scheme consisting of bacterial species as the rows, and the scores from the computational or validation outcomes as columns. Cohen’s kappa was then calculated for each table using the “kappa2” function from the “irr” R package (https://CRAN.R-project.org/package=irr).

## Data Availability

This project used previously acquired publicly available data(20). All code that was written for this project can be found in the GitHub repository: https://github.com/KarstensLab/BladderBacteriaSpecies.

## Acknowledgements

This work was supported by the National Institutes of Health (NIH): NIDDK award number K01 DK116706 (LK); NCAT award number UL1TR002369 (LK); NIA award numbers R03 AG060082 and P30AG028716 (NS). The content of the manuscript is solely our responsibility and does not represent the official views of the NIH or any other funding agency

## Author contributions

CH and LK conceived and designed experiments, and wrote the manuscript. CH performed analyses, LK reviewed analyses. AJW supplied the data. LK, CH, MM, HS, TG, IF, NS, and AJW interpreted results and revised the manuscript.

## Supplemental Figures

**Supplemental Figure 1.**
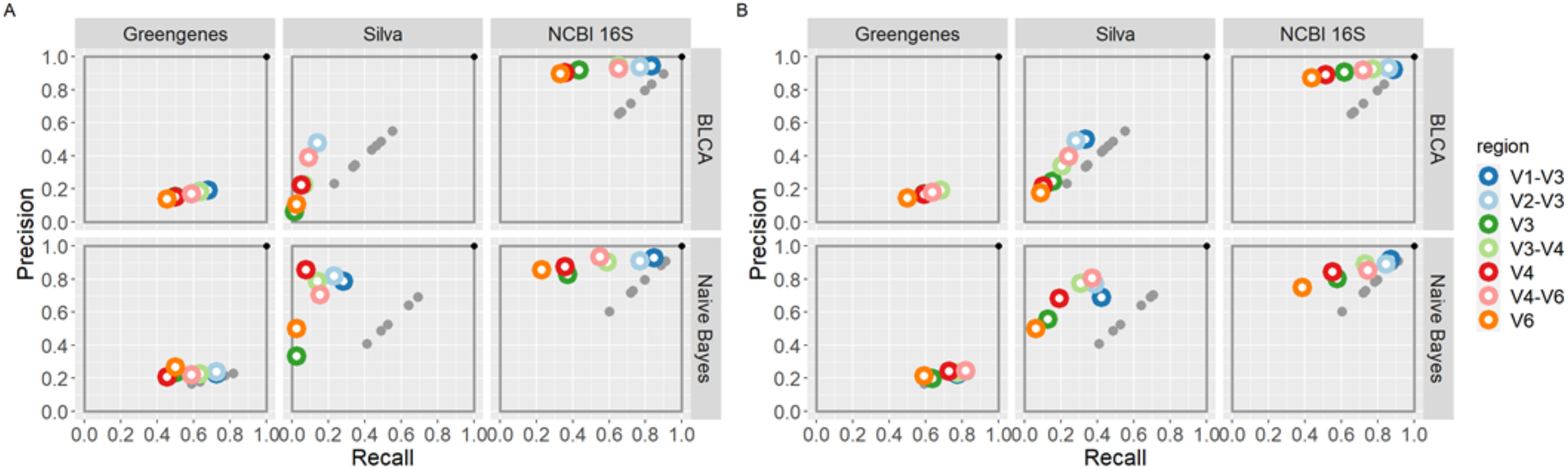
Classification scheme precision and recall when using a confidence scores of (A) 80% and (B) 50% as a threshold. The schemes that use the Silva database have very low recall compared to when the confidence score is ignored (gray dots), whereas schemes that use Greengenes and NCBI 16S are not as affected.

**Supplemental Figure 2.**
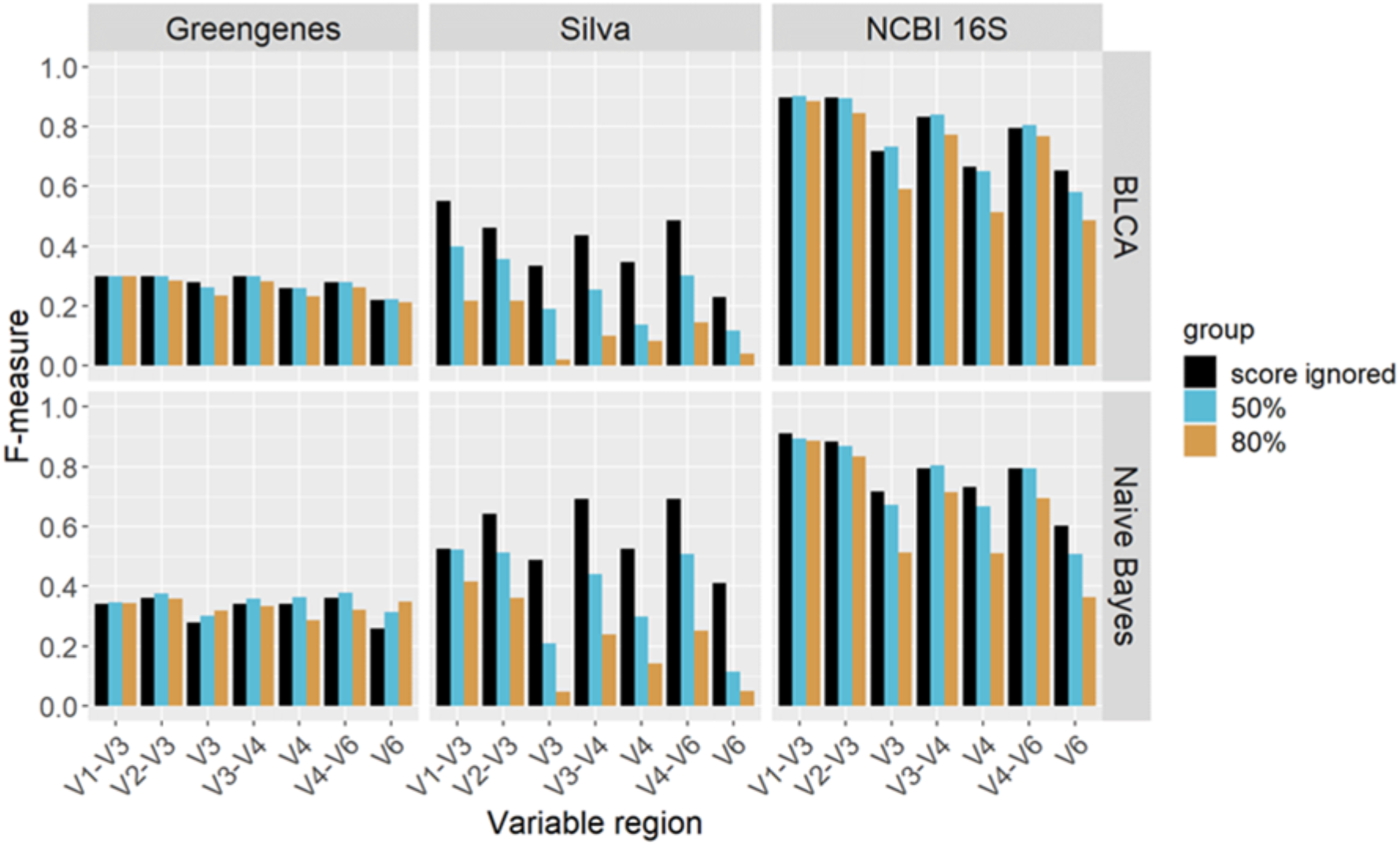
F-measure values for all classification schemes. Values shown are for confidence scores of 50%, 80%, and when confidence scores are ignored.

**Supplemental Figure 3.**
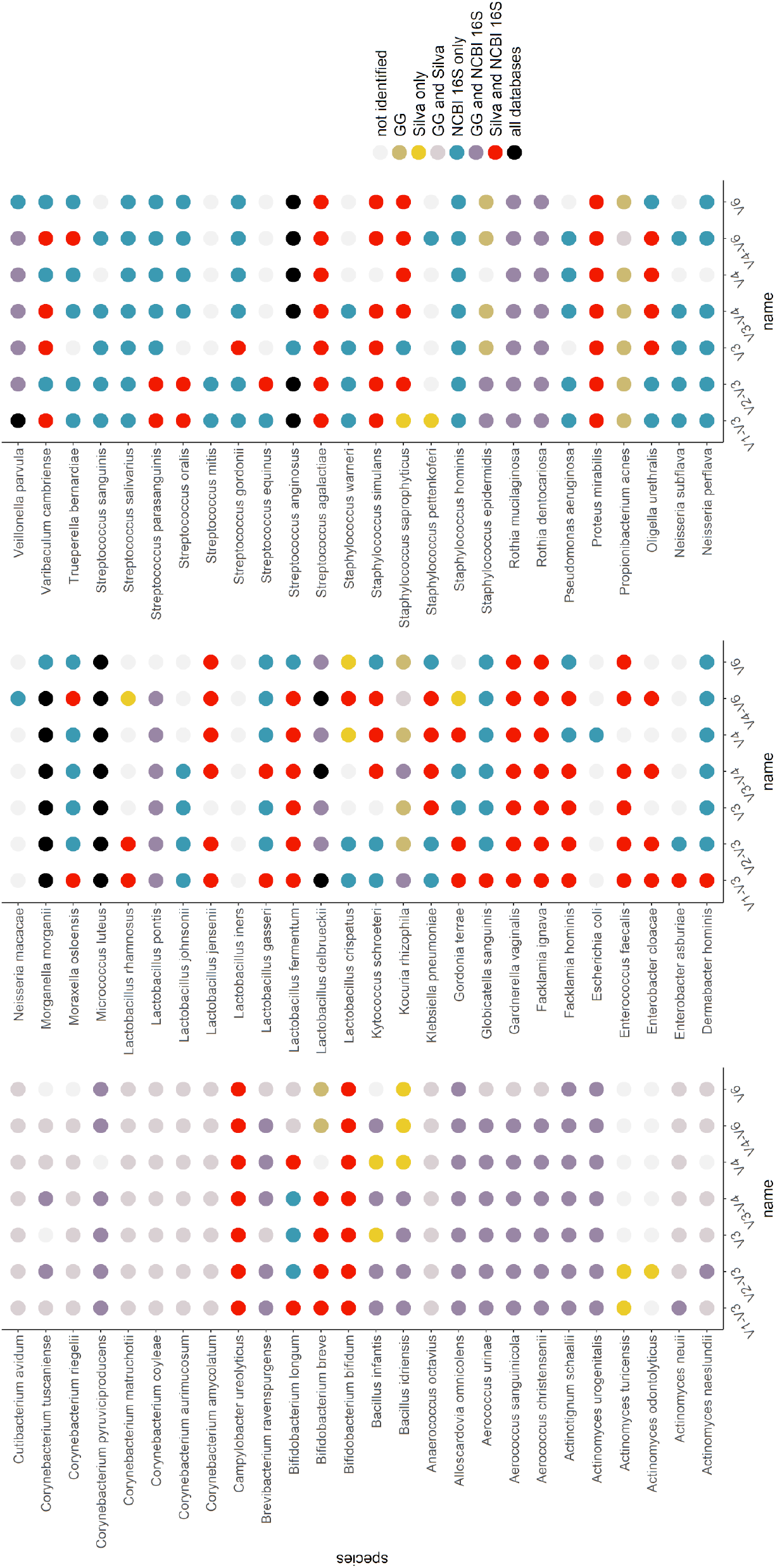
Bladder bacterial species of the Thomas-White dataset identified by database, variable region, and BLCA classifier.

**Supplemental Figure 4.**
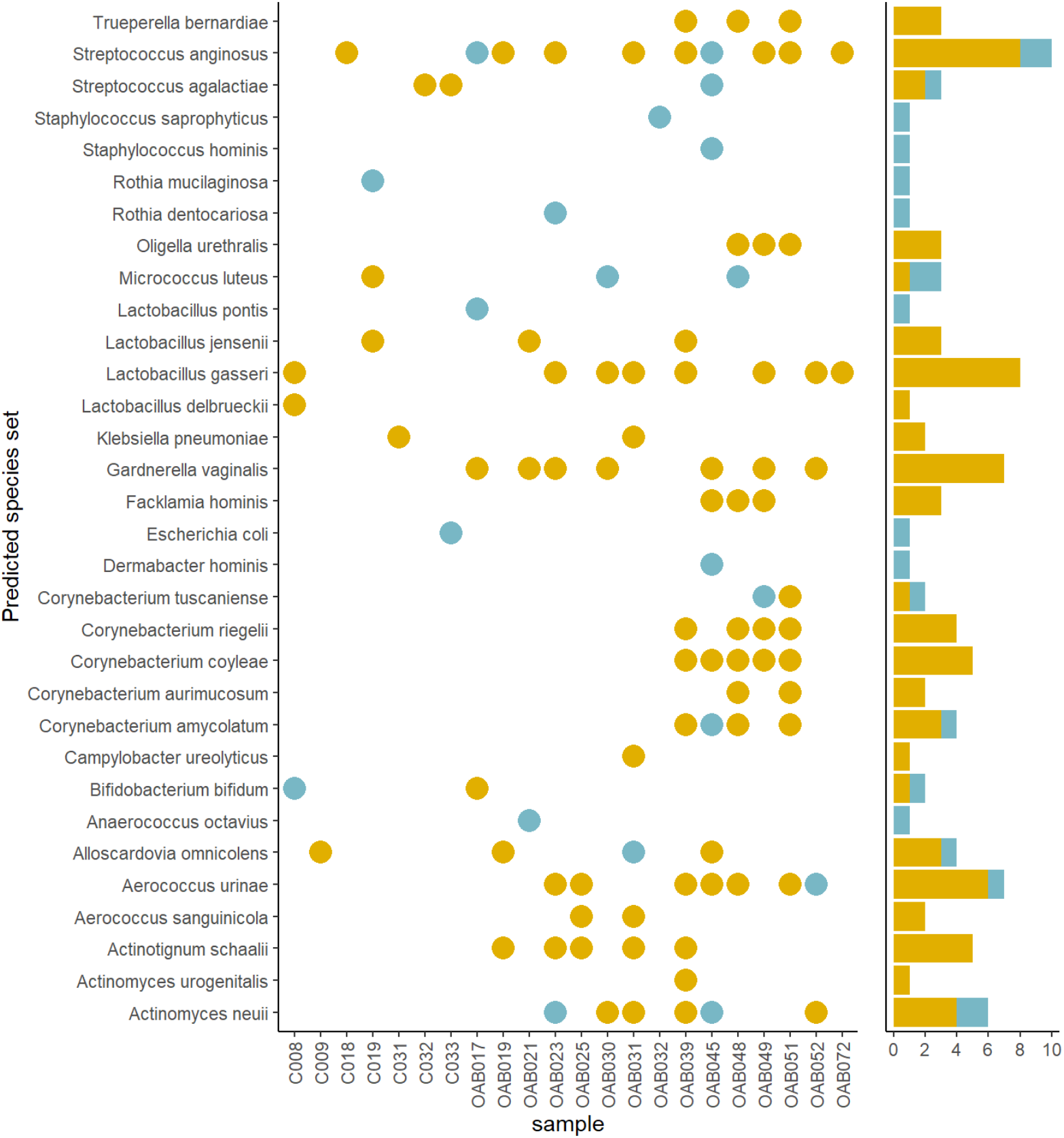
Summary of the NCBI 16S classification for *in silico* classification on the V4 validation dataset. Blue dots represent the species/sample pair that was identified by whole genome sequencing, but was not identified by the classification scheme. Yellow dots represent the species/sample pair that was successfully identified by the classification scheme and whole genome sequencing of the culture data.

